# Effects of flavorless electronic cigarette aerosol on the survival and growth of common oral commensal streptococci

**DOI:** 10.1101/529909

**Authors:** Giancarlo A. Cuadra, Maxwell Smith, John M. Nelson, Emma Loh, Dominic Palazzolo

## Abstract

The use of electronic cigarettes (ECIG) has become very common. Consequently, critical analysis of the biological effects of ECIG aerosol deserves attention. Since the mouth is the first anatomical cavity exposed to aerosol, the oral bacteria within are also exposed. We hypothesize that while cigarette smoke has a detrimental effect on the survival and growth of oral commensal streptococci, flavorless ECIG aerosol does not. Survival and growth of several strains of commensal streptococci were measured after exposure to flavorless ECIG aerosol ± nicotine and smoke. Peristaltic pumps were used to transport flavorless aerosol ± nicotine or cigarette smoke into chambers containing recently seeded colony forming units of four strains of oral commensal streptococci on agar plates. Bacterial survival and growth, based on colony counts and sizes, were determined 24 hours post-exposure. Lastly, aerosol or smoke were delivered into chambers containing the four strains of streptococci pre-adhered to plastic coverslips. Bacterial survival and growth, as indicated by biofilm formation, were determined 24 hours post-exposure via scanning electron microscopy. The results suggest that flavorless aerosol ± nicotine has a modest effect on bacterial growth both as colonies on agar and as biofilms. In contrast, smoke dramatically decrease bacterial survival and growth in all parameters measured. Therefore, unlike cigarette smoke, flavorless ECIG aerosol has only a small effect on the survival and growth of oral commensal streptococci.

## Introduction

The use of electronic cigarettes (ECIG), referred to as vaping, has gained immense popularity in recent times [1]. Cigarette smoke is known to contain thousands of detrimental compounds, but the constituents of flavorless ECIG aerosol are few. In general, ECIG-liquid (E-liquid) consists of propylene glycol and/or vegetable glycerin, nicotine ranging from 0 to >24 mg/ml and a variety of flavors [2]. While vaping on ECIG devices is commonplace, as of yet there is no clear evidence of the potential issues its usage could cause. For this reason, the physiological effects of ECIG aerosol should be seriously investigated.

Currently, there are few studies regarding effects of ECIG-generated aerosol on physiological systems as compared to cigarette smoke. A few reports claim that ECIG use is as dangerous (or more dangerous) than traditional smoking [3–5]. E-liquid flavorings have also recently been reported to induce inflammatory and oxidative responses in human monocytic cell lines [6]. Similarly, various flavored E-liquids have a toxic effect on stem cells and terminally differentiated cell lines [7]. Moreover, human bronchial epithelial as well as oral epithelial cell lines exposed to ECIG-generated aerosol with flavorings increased pro-inflammatory cytokine production and caused other adverse effects on the biology of these cell models [8–10]. All these studies indicate that ECIG-generated aerosol containing flavorings can be detrimental to exposed tissues and therefore deserves more attention and information to the public.

The oral cavity contains a vast diversity of commensal, opportunistic and sometimes pathogenic bacteria. The most common types of commensal bacteria are streptococci [11,12], which are found in individuals at any level of oral health and disease [13,14]. Among these bacteria, some of the most common species are *Streptococcus gordonii, Streptococcus intermedius, Streptococcus mitis* and *Streptococcus oralis* [15–17]. All four of these species are crucial in the development of oral biofilms on both soft and hard surfaces within the mouth [13,18,19]. These species are considered commensal early colonizers [20–22]. All four species are beneficial to the host oral cavity in the context of their interactions with pathogenic species related both to caries and periodontal disease [23–28]. By extension, since oral health and overall general health are directly correlated, any disruption to the bacterial flora within the oral cavity could lead to systemic disease, especially certain types of cardiovascular disease [29]. For this reason, it is important to examine how vaping affects the oral microbiota.

Smoking has been reported to be a leading risk factor for caries and periodontal disease [30–33] and is known to considerably affect the subgingival oral microbiome *in situ* [34]. No studies (to our knowledge) are available to show how ECIG aerosol specifically affects oral commensal streptococci known to provide a protective barrier against external insults. Since *S. gordonii*, *S. intermedius*, *S. mitis* and *S. oralis* are crucial in the development of oral biofilms on both soft and hard surfaces within the mouth, the aim of this work is to test the impact of flavorless ECIG aerosol and compare it to conventional cigarette smoke on the survival and growth of oral commensal streptococci. Our findings seem to demonstrate that flavorless ECIG aerosol has little to no effect on the survival and growth of commensal oral streptococci.

## Materials and methods

### Reagents and supplies

All reagents and supplies for conducting these investigations were purchased from Thermo Fisher Scientific (Waltham, MA) unless otherwise noted.

### Bacterial strains

*S. gordonii* DL1, *S. intermedius* 0809, *S. mitis* UF2 and *S. oralis* SK139 were kindly provided by Dr. Robert Burne from the University of Florida. All strains were grown in brain heart infusion (BHI) broth with 5 μg/ml hemin or BHI agar at 37°C and 5% CO_2_. Bacterial stocks were stored at −80°C.

### E-Liquid

E-liquid was composed of 50% propylene glycol and 50% vegetable glycerin (*i.e*. glycerol) with or without (±) 20 mg/ml of (S)-(-)-nicotine (Alpha Aesar, Tewksbury, MA). No flavors were added. This nicotine concentration on a per cigarette equivalent is higher than the typical concentration of nicotine in a tobacco cigarette [35].

### Exposure apparatus

Bacterial samples were exposed to either air, flavorless ECIG aerosol ± 20 mg/ml nicotine or cigarette smoke following already established protocols [36,37]. Briefly, Cole-Palmer Master Flex L/S peristaltic pumps (Vernon Hills, IL) and tubing were used to simulate puffing and transport air, smoke or aerosol into an acrylic chamber, as shown in Figure 1. Peristalsis and flow rates were adjusted to 400 ml/min or 33.3 ml in five seconds as indicated in Table 1. Puffing was conducted at 5 seconds on (pumps active) followed by a ten-second rest period (pumps inactive). The puffing protocol consisted of 0, 10, 25, 50 and 75 puffs. Values of total nicotine exposure in the acrylic chamber are shown in Table 1. All pump-puffing experiments were conducted within a P20 Purair ductless fume hood (Airscience, Fort Myers, FL) with a HEPA filter.

**Fig 1.**
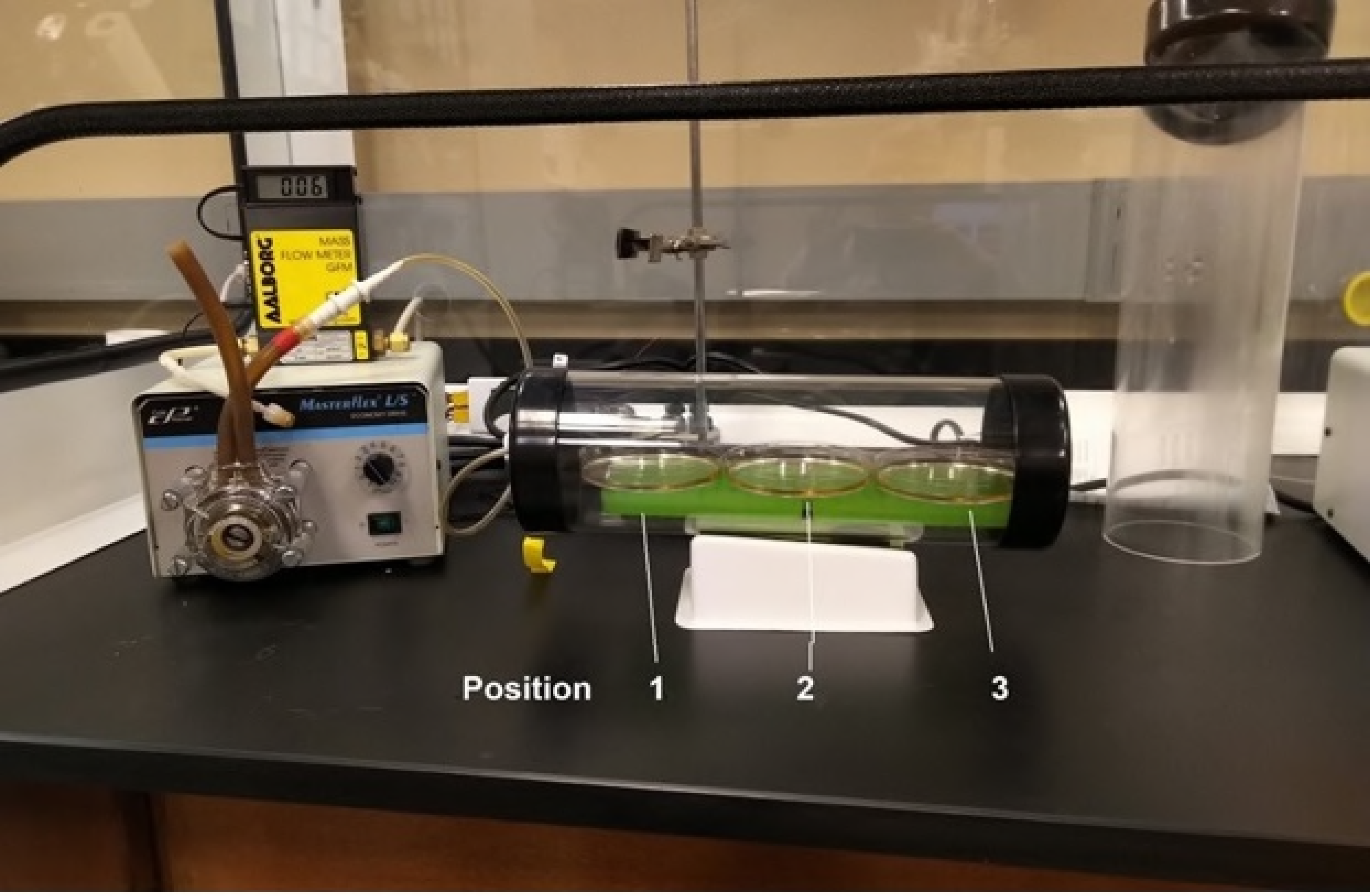
Exposure apparatus for flavorless ECIG aerosol and cigarette smoke. Peristaltic pumps, tubing and acrylic chamber containing three plates numbered 1, 2 and 3 according to their relative position with respect to the aerosol or smoke source (tubing connection to chamber). Mass flow meter (small yellow and black machine) was used to calibrate the flow of materials at 400 ml/min. All materials are shown inside a Purair ductless fume hood.

**Table 1:** Pump/chamber parameters and nicotine concentrations within the exposure chambers.

|  | **Smoke**<br>**pump/chamber** | **Aerosol**<br>**pump/chamber** |
| --- | --- | --- |
| **Pump Flow Rate (ml/min)** | 400 | 400 |
| **Puff Duration (seconds)** | 5 | 5 |
| **Puff Volume (ml)** | 33.3 | 33.3 |
| **Nicotine (mg/cigarette)<sup>a</sup>** | 0.92 | 2.80 |
| **Nicotine (mg/puff)<sup>b</sup>** | 0.06 | 0.19 |
| **Nicotine Flow (µg/puff/ml) into Chamber** | 1.8 | 5.7 |
| **Exposure Chamber Volume (cm<sup>3</sup>)** | 2126 | 2126 |
| **Nicotine Delivered to Chamber (mg)** |  |  |
| 0 puffs (0.0 cigarettes) | 0.0 | 0.0 |
| 25 puffs (1.7 cigarettes) | 1.5 | 4.8 |
| 50 puffs (3.3 cigarettes) | 3.0 | 9.5 |
| 75 puffs (5.0 cigarettes) | 4.5 | 14.3 |
| **Nicotine Concentration in Chamber (µg/cm<sup>3</sup>)<sup>c</sup>** |  |  |
| 0 puffs (0.0 cigarettes) | 0.0 | 0.0 |
| 25 puffs (1.7 cigarettes) | 0.7 | 2.2 |
| 50 puffs (3.3 cigarettes) | 1.4 | 4.5 |
| 75 puffs (5.0 cigarettes) | 2.1 | 6.7 |

<sup>a</sup>For smoke, the value of mg nicotine per Marlboro® Red cigarette [35]. For aerosol, the value 2.8 mg/cigarette is based on 9.3 µl of E-liquid (20 mg/ml nicotine) aerosolized per puff and that 15 puffs is equivalent to one cigarette [40].
<sup>b</sup>One cigarette is equivalent to 15 puffs.
<sup>c</sup>These values assume that all nicotine remains within the chamber, which is not the case since the exposure chambers are fitted with rubber end caps perforated with small holes to allow venting of the exposure chambers [36,37].

### Distribution of aerosol and smoke in the exposure chamber

Three 100 mm plates containing 10 ml of BHI broth were placed in the exposure chamber. According to Figure 1, the plate in position 1 is the closest to the source of aerosol or smoke and the plate in position 3 is the furthest away. BHI broth in three plates was exposed to flavorless ECIG aerosol with 20 mg/ml nicotine and cigarette smoke for a total of 10, 25, 50 and 75 puffs following above protocols. Concentrations of nicotine in the BHI broth of plate 1, plate 2 and plate 3 (Figure 1) were evaluated by high performance liquid chromatography (HPLC).

### HPLC determination of nicotine

Standard solutions of 99% (S)-(-)-nicotine, were prepared in BHI broth at concentrations of 0.4, 0.2 and 0.1 mg/ml. Standards and samples of BHI exposed to 10, 25, 50 or 75 puffs of flavorless ECIG aerosol with nicotine or conventional cigarette smoke were analyzed by HPLC coupled with photodiode array detection as previously described [38,39]. A Shimadzu HPLC system (Columbia, MD) was used to quantitate nicotine and included the following: a photodiode array detector (SPD-M20A), dual pumps (LC-20AT), a column oven (CTO-20A), an in-line membrane degasser (DGU-20A3R) and a Rheodyne 7725I manual injector with 20 µl loop (40 µl injection volume). Nicotine was separated on a Phenomenex (Torrance, CA) 15-cm, Kinetex^®^ 5µm reversed phase C-18 column preceded by a Phenomenex Security Guard. Column temperature was maintained at 35°C. Nicotine was detected at UV wavelengths between 230 and 300 nm and quantifications were carried out at 260 nm. The mobile phase was delivered at a rate of 1 ml/minute in gradient fashion where mobile phase A consisted of 10% acetonitrile in 20 mM ammonium formate adjusted to pH 8.5 with 50% ammonium hydroxide and mobile phase B consisted of 100% acetonitrile. Mobile phase A decreased from 100% to 80% from 0 to 10 minutes, decreased from 80% to 20% from 10 to 20 minutes, increased from 20% to 100% from 20 to 21 minutes and remained at 100% till the end of the run time at 30 minutes. Mobile phase B increased from 0% to 20% from 0 to 10 minutes, increased from 20% to 80% from 10 to 20 minutes, decreased from 80% to 0% from 20 to 21 minutes and remained at 0% till the end of the run time at 30 minutes. The nicotine standard curve was linear (R^2^ = 0.9998) and nicotine eluted at a retention time of 10.5 minutes. Chromatographic parameters were PC-controlled using a Lab Solutions work station.

### Colony forming units (CFU)

Starter overnight cultures of all four strains of bacteria were adjusted to OD 595 nm of 1.0 and serially diluted to numbers permissible for CFU counting. Twenty microliters of each species were plated in triplicates on BHI agar plates. As soon as the 20 µl volume dried into the agar, bacteria were exposed uncovered to air, flavorless ECIG aerosol (± 20 mg/ml nicotine) or cigarette smoke for up to 75 puffs. Following exposures, plates were incubated at 37°C and 5% CO_2_ for 24 hours. The next day, colonies were digitally photographed using a Moticam 1080 HDMI & USB camera (Motic®, Richmond, British Columbia, Canada) attached to a Fisher brand stereomicroscope and counted. Average colony sizes (as indexed by the area of individual colonies) were determined using the Moticam supplied on-board camera software.

### Biofilm Biomass

Starter overnight cultures of bacteria were adjusted to OD 595 nm of 1.00. After adjustment, 100 μl of each culture was seeded separately on sterile plastic coverslips (13 mm diameter) in 12-well plates. Bacteria were allowed to adhere to the surface of the coverslips for 1 hour at 37°C, 5% CO_2_ and the excess unbound bacteria were washed 3 times with 0.5 ml PBS. Excess liquid on the coverslips was removed and the 12-well plates containing the coverslips were exposed uncovered to air, flavorless ECIG-generated (± 20 mg/ml nicotine) or cigarette smoke for up to 75 puffs. Following exposure, 1 ml of 50% BHI broth (v/v in sterile water) was added to each well of the 12-well plate ensuring that exposed coverslips were completely submerged. Exposed bacteria were subsequently incubated for 24 hours at 37°C, 5% CO_2_ to allow for biofilm growth on the coverslips. At the end of the 24-hour incubation period, BHI broth was removed from the wells and the coverslips were washed 3 times with 1 ml PBS to remove excess unbound bacteria. Biofilms were fixed with 1 ml of 4% formaldehyde for at least 30 minutes. Coverslips were then processed for SEM imaging (described below).

### Biofilm processing and SEM imaging

The 4% formaldehyde was removed from each well and each coverslip was rinsed two times with 1 ml of deionized water. The biofilms on the coverslips were then dehydrated using an increasing alcohol gradient (*i.e*. 30 minutes in each of 50, 70, 90 and 100% ethanol) followed by chemical drying with 98% hexamethyldisilizane for 30 minutes. The coverslips with attached biofilms were then removed from the 12-well plates and air dried for 5 to 10 minutes before mounting onto 13 mm aluminum pin-type stubs [Structure Probe, Inc. (SPI), West Chester, PA]. Conductive, 12 mm diameter, double-sided carbon-impregnated adhesive disks (SPI) were used to adhere the coverslips to the stubs and 1 to 2 hours was allowed for complete adherence. In the mounting process, extreme care was used to ensure the side of the coverslip with the bacterial biofilm was facing up and not disrupted. The mounted bacterial biofilms were then sputter coated using a Hummer IV-A sputtering system (Anatech Ltd., Alexandria, VA) and plated with 300Å of 1:1 gold:palladium. SEM images of biofilms grown on coverslips were taken with a TOPCON ABT-60 microscope at an acceleration voltage of 15 kV and a magnification of 450X.

### Statistical analysis

Mean and standard error of the mean (SE) were calculated for nicotine in BHI broth. One-way ANOVA followed by Newman-Keuls multiple comparison test analysis was used to determine differences in nicotine concentrations between plate positions 1, 2 and 3 after 10, 25, 50 and 75 puffs of flavorless ECIG aerosol with 20 mg/ml nicotine or conventional cigarette smoke. CFUs were visually counted and the average of three largest colonies in each quadrant of an agar plate were used to determine mean colony size for all bacteria at every exposure. Mean and SE were calculated for CFU counts and colony size. Statistical variance between groups was determined using a two-way ANOVA, followed by Bonferoni post hoc analysis. Differences were considered statistically significant when p < 0.01.

## Results

### Distribution of flavorless ECIG aerosol and smoke in the exposure chamber

Figure 1 shows the setup of three plates in tandem inside the acrylic chamber. The BHI broth in all three plate positions received comparable amounts of nicotine (Fig 2, p > 0.05). The results also show that the amount of nicotine, regardless of source, increases in a puff-dependent manner in all three plates. Lastly, the results also show the projected results of higher levels of nicotine from flavorless ECIG aerosol compared to cigarette smoke (Fig 2) and agree with the expected values shown in Table 1. These data indicate that plate position is not a confounding factor in the results and interpretation of the following experiments.

**Fig 2.**
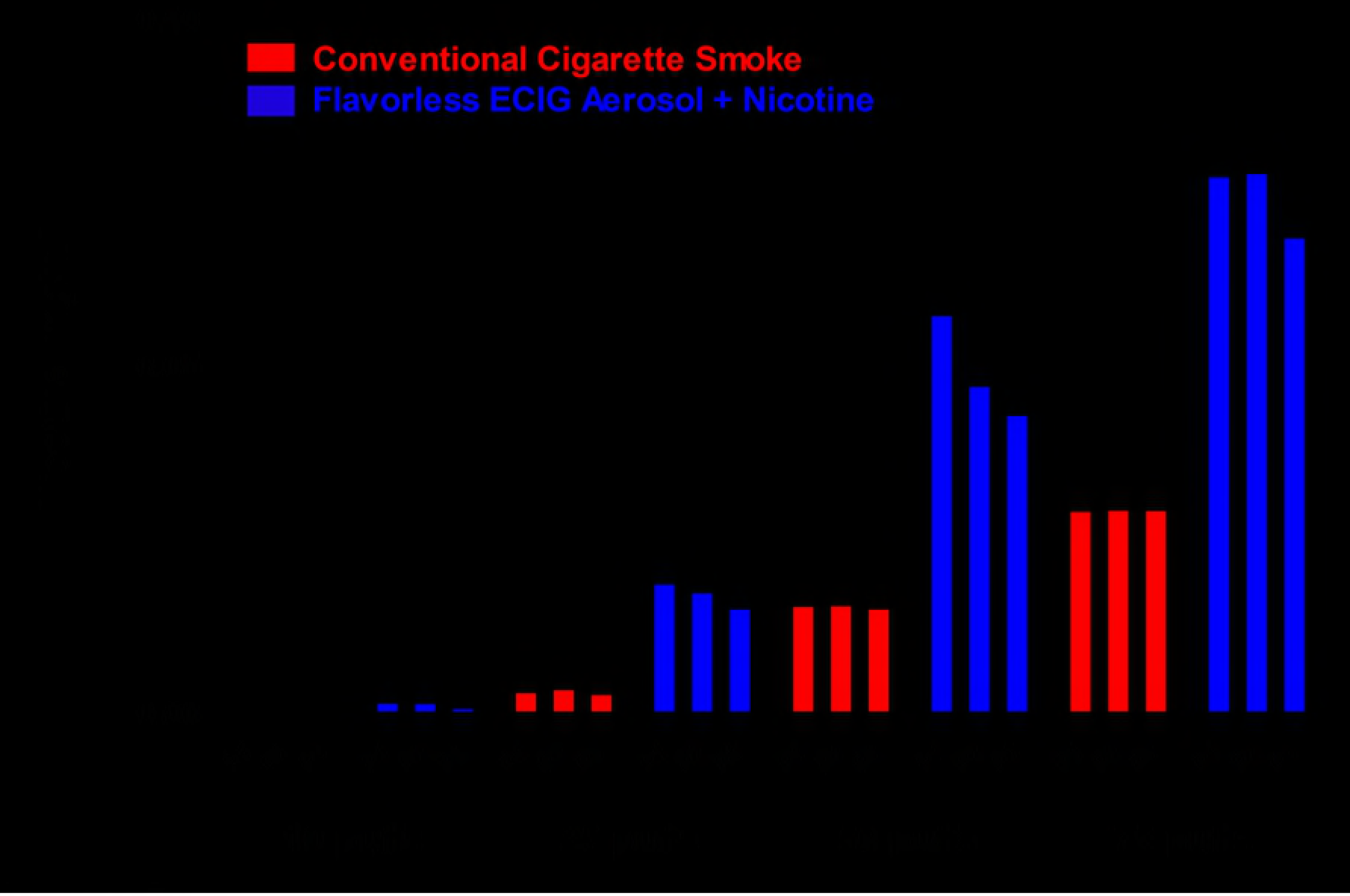
Levels of nicotine in three plates inside the exposure chamber. Nicotine concentrations (mg/ml) in three plates of BHI broth (*i.e*. P1, P2 and P3) set up in tandem inside an exposure chamber after 10, 25, 50 and 75 puffs of flavorless ECIG aerosol containing 20 mg/ml nicotine or conventional cigarette smoke. Each data point is the Mean ± SE (n=3).

### Effects of flavorless ECIG aerosol and smoke on CFU counts

CFU counts of commensal oral streptococci seeded on agar and exposed to puffs of air (control), flavorless ECIG aerosol ± nicotine and cigarette smoke prior to overnight colony growth are shown in Figure 3. Without exposure (0 puffs), the number of CFUs per agar plate ranged between 37 and 63 for *S. gordonii*, 25 and 42 for *S. intermedius*, 35 and 70 for *S. mitis* and 65 and 84 for *S. oralis*. Bacteria exposed to flavorless ECIG aerosol ± nicotine grow similar numbers of colonies as compared to those exposed to air, although significant differences (p < 0.01) between aerosol with and aerosol without nicotine exist for *S. gordonii* and *S. mitis* at 75 and 50 puffs, respectively (Fig 3). In drastic contrast, bacteria exposed to 50 or 75 puffs of cigarette smoke yield no colonies at all. Our results indicate a profound toxic effect of cigarette smoke and a far lower toxic effect of flavorless ECIG aerosol on commensal oral streptococci.

**Fig 3.**
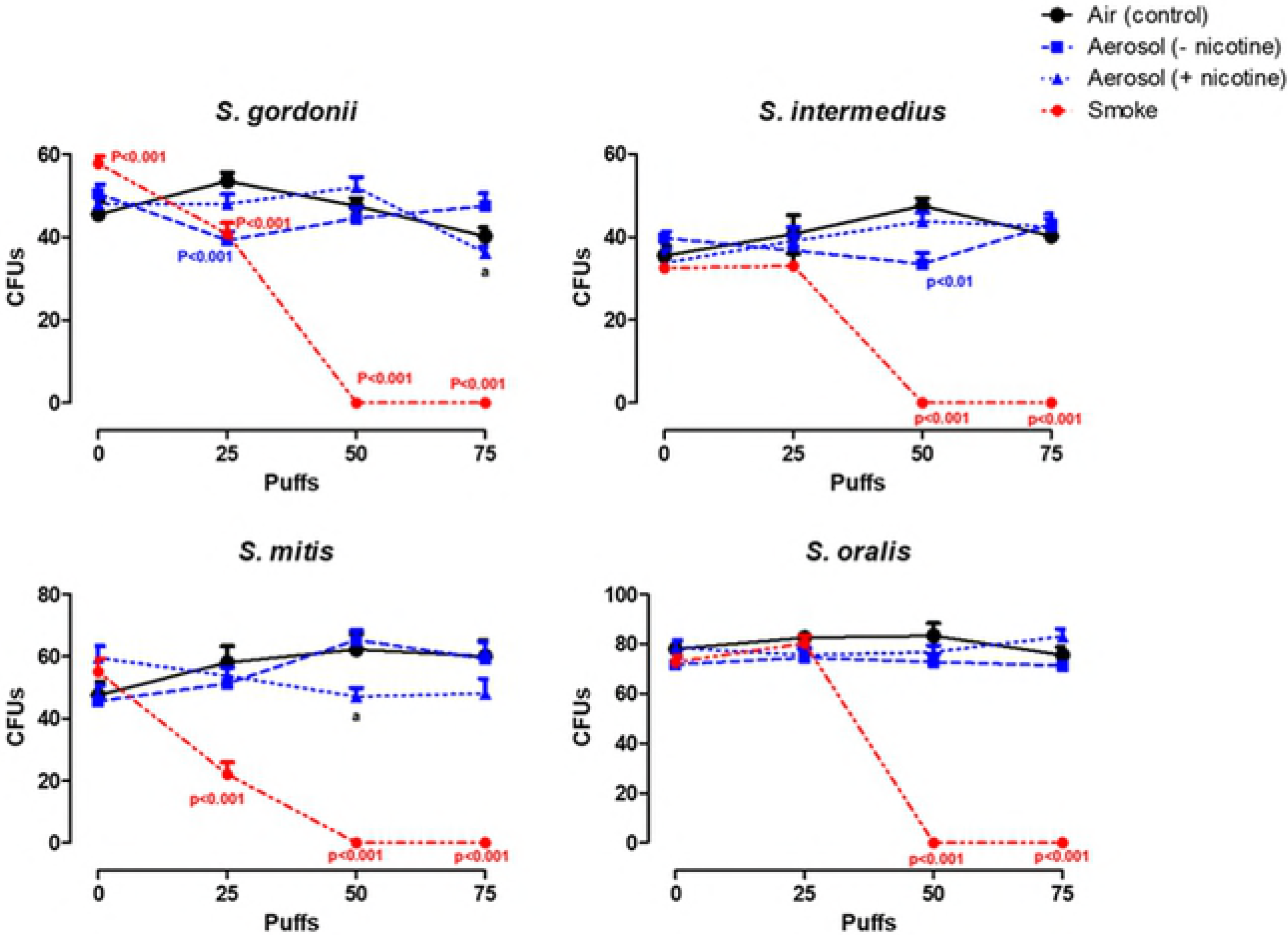
Effect of air, flavorless ECIG aerosol (± 20 mg/ml nicotine), or smoke (Marlboro^®^ Red cigarette) on CFU counts. Each data point is Mean ± SE, n=4 (average of triplicates from each quadrant of an agar plate), p values indicate significance from control. a = p < 0.01 between aerosol with nicotine and without nicotine.

### Effects of flavorless ECIG aerosol and smoke on colony size

Besides the obvious absence of colonies following exposure to 50 and 75 puffs of smoke, colonies exposed to 25 puffs of cigarette smoke also appear to have a smaller size compared to those colonies exposed to air or flavorless ECIG aerosol ± nicotine. Figure 4A displays the average area of colony sizes for *S. gordonii*, *S. intermedius*, *S. mitis* and *S. oralis* without exposure (0 puffs) which are 0.543 ± 0.023, 0.244 ± 0.015, 0.339 ± 0.018 and 0.110 ± 0.003 mm^2^, respectively. Figure 4B highlights the smaller sizes of colonies for all bacteria after exposure to 25 puffs of smoke compared to colonies after exposure to zero puffs. Colony sizes after exposure to 0, 25, 50 and 75 puffs of air (control), flavorless ECIG aerosol ± nicotine and cigarette smoke are quantified in Figure 4C. The average colony size of all bacteria exposed to 25 puffs of smoke are significantly smaller (p < 0.01 to p < 0.001) than the controls. Flavorless ECIG aerosol without nicotine also appears to have a slight effect (p < 0.01) on the colony size of *S. gordonii* after exposure to 50 puffs.

**Fig 4.**
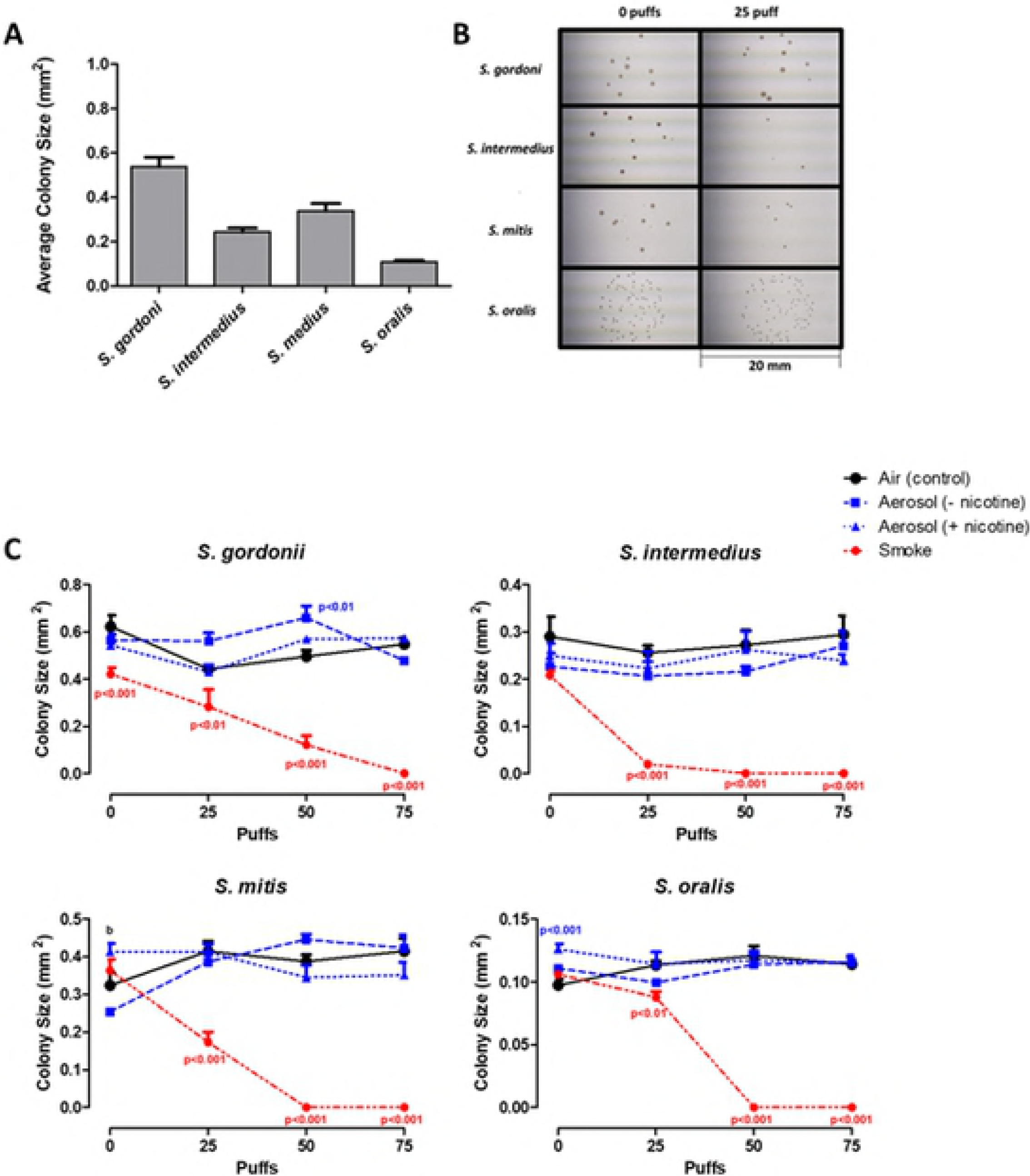
Effect of air, flavorless ECIG aerosol (± 20 mg/ml nicotine), or smoke (Marlboro^®^ Red cigarette) on CFU size. A. Average colony size of commensal oral streptococci. The average of the three largest colonies in each quadrant of four agar plate (n=16) were used to determine mean colony size for all bacteria at every exposure. Each data point is the Mean ± SE (mm^2^). **B.** Representative images of colony sizes of commensal oral streptococci following exposure of 0 and 25 puffs of smoke (Marlboro^®^ Red cigarette). Each frame is 20 mm long. **C.** Effect of air (control), aerosol (± 20 mg/ml nicotine), or smoke (Marlboro^®^ Red cigarette) on colony size of commensal oral streptococci. Each data point is Mean ± SE, n=4, p values indicate significance from control. b = p < 0.001 between aerosol with nicotine and without nicotine.

### Effects of flavorless ECIG aerosol and smoke on bacterial biofilms

Figure 5 illustrates the formation of single-species bacterial biofilms on plastic coverslips 24 hours after exposure to 0 puffs of air (control), 75 puffs of air (control), flavorless ECIG aerosol ± nicotine and cigarette smoke. As shown, each of these species is able to grow biofilms after exposure to air (control) and flavorless ECIG aerosol ± nicotine, but not cigarette smoke. Compared to air exposures (0 and 75 puffs), 75 puffs of flavorless ECIG aerosol ± nicotine is permissive for biofilm formation and growth regardless of the overall architecture of bacterial communities for all four species. This indicates that flavorless ECIG aerosol ± nicotine has little to no effect on oral commensal streptococci biofilm formation and growth under these conditions.

**Fig 5.**
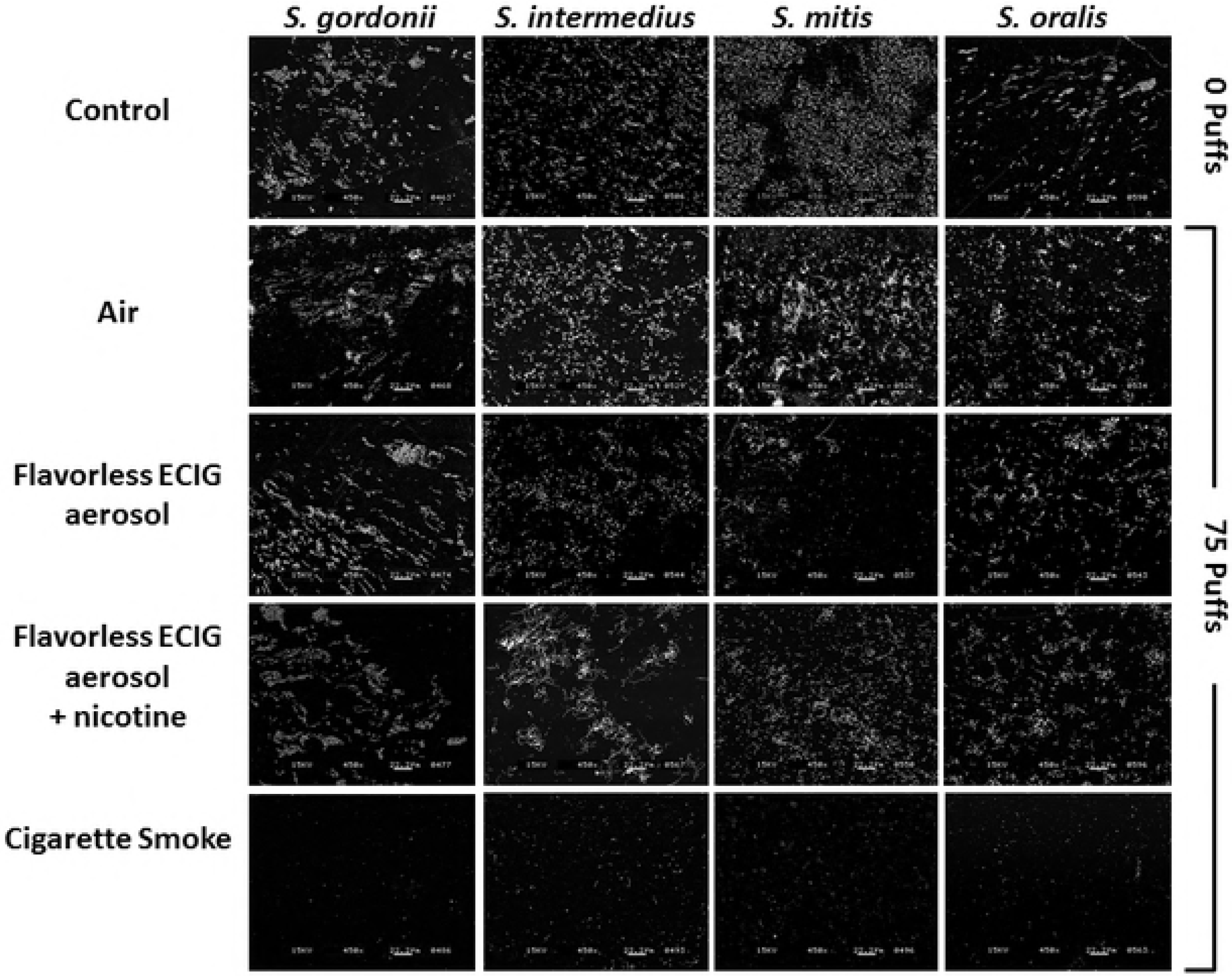
Effect of air, flavorless ECIG aerosol (± 20 mg/ml nicotine), or smoke (Marlboro^®^ Red cigarette) on biofilm formation. Representative images of biofilm formation for commensal oral streptococci following exposure to 0 puffs of air (control) or 75 puffs of air (control), aerosol (± 20 mg/ml nicotine) and smoke (Marlboro^®^ Red cigarette). All images were acquired at 450X using an acceleration voltage of 15 kV.

## Discussion

The current work demonstrates that flavorless ECIG aerosol ± nicotine has little to no toxic effect on the *in vitro* growth of the four oral commensal streptococci tested here. Our data show that CFUs for all four species exposed to flavorless ECIG aerosol ± nicotine can grow similar numbers and to similar sizes as compared to their untreated counterparts. Our data also demonstrate that bacteria attached to coverslips and exposed to flavorless ECIG aerosol ± nicotine are also able to grow biofilms like their untreated controls. However, when bacteria are exposed to cigarette smoke, growth of colonies and biofilms is severely impaired or completely obliterated. Furthermore, it is evident that nicotine is not the culprit of this impairment since ECIG aerosol contained a higher concentration of nicotine than cigarette smoke on a per puff basis (Figure 2). Altogether, based on these results we propose that flavorless ECIG aerosol ± nicotine does no apparent harm to these four oral bacteria under the conditions tested.

Overall, flavorless ECIG aerosol (± nicotine) appears to have little or no effect on CFU number and colony size of all bacteria tested (Figures 3 and 4). *S. gordonii* and *S. intermedius* exhibit a slight decrease in CFU number when exposed to 25 and 50 puffs of aerosol without nicotine, respectively, and *S. gordonii* exhibits a slight increase in colony size when exposed to 50 puffs of aerosol without nicotine. Likely, these differences are due to inherent variability associated with CFU plating [41] and not an effect contradictory to the overall trend. Furthermore, the presence or absence of nicotine in the flavorless ECIG aerosol also appears to have little effect on CFU number and colony size of all streptococci tested. The results of this investigation are comparable to the results of Huang *et al*. (2014) who report no significant difference in planktonic growth of *S. gordonii* in tripticase soy broth (TSB) growth media or of CFU counts on TSB agar plates at nicotine concentrations below 1 mg/ml [31], although nicotine concentrations between 1 and 4 mg/ml appeared to stimulate *S. gordonii* planktonic growth in a dose-dependent manner. Most likely the effect of nicotine on streptococcal bacteria is species-dependent, since Li *et al*. (2014) determined that nicotine had little effect on *S. sanguinis* biofilm formation, but increased *S. mutans* biofilm formation [30].

Oral bacteria live in polymicrobial communities [42], even when exposed to cigarette smoke. The study by Shah *et al*. (2017) indicates that, in polymicrobial biofilms, commensal species typically suffer and struggle to grow in the presence of tobacco smoke while pathogenic species thrive under the same conditions [43]. The decrease in growth of commensal bacteria exposed to cigarette smoke (Figs 3, 4 and 5) may be due to downregulation of important metabolic factors in the commensal species [43]. Interestingly, Zonuz et al., (2008) reported accelerated growth of *S. sanguinis* and *S. mutans* in the vicinity of cigarette smoke [44] - an apparently contradictory finding. The findings of the present study are in agreement with Shah *et al*. (2017), in which commensal bacteria are unable to form single-species biofilms when exposed to cigarette smoke. Assuming the microbial landscape within the oral cavity shifts toward poor oral health in response to cigarette smoke, as Shah *et al*. (2017) indicate, this could ultimately lead to more severe pathophysiologic problems [43]. For example, the report by Bagaitkar *et al*. (2011) suggests that cigarette smoke extract augments the persistence of *P. gingivalis* in biofilms with *S. gordonii* by elevated expression of major fimbrial antigen [33]. *P. gingivalis* is a Gram-negative pathogenic bacterium and a principle periodontitis inducing agent [45,46]. Periodontal pathogens induce systemic inflammation, ultimately leading to increased risk of cardiovascular disease such as atherosclerosis [47,48]. While floverless ECIG aerosol has no effect on our commensal bacteria’s ability to form biofilms (Figure 5), it is important to test the effects of conventional cigarette smoke and ECIG aerosol with mixed-species biofilms to determine the effects of the same environmental agents in an open system. The results of such experiments will give a much better understanding of the effects of smoke and ECIG aerosol on oral microbial communities. Our experimental design also tested only one strain of bacteria at a time. Interactions between strains of commensal bacteria, as well as with pathogenic bacteria, should be investigated to ultimately determine if the effect of flavorless ECIG-aerosol is also different than that of smoke in mixed species biofilms. To best address this question, the bacteria should be cultured in an open system following exposure. An open system design would resemble the natural oral environment in the context of mixtures of oral microbiota combined with salivary flow, which aids in clearance of external materials from the oral cavity.

As CFU counts, colony size and biofilm formation obtained from this investigation indicate, we are confident that flavorless ECIG aerosol has a less drastic effect on the oral commensal bacteria tested than cigarette smoke. However, this study does have limitations. The in-house prepared E-liquid used in this study represents a single rendition of E-liquid and does not represent the flavored preferences of most ECIG users. In addition, the only concentration of nicotine used is 20 mg/ml and the only ratio of propylene glycol to vegetable glycerin is 1:1 v/v. Therefore, conclusions concerning the effects of ECIG aerosol on the survival and growth of commensal bacteria must be considered within the context of variability in the composition of commercially available E-liquids. Another limitation is the fact that flavorless ECIG-aerosol and conventional cigarette smoke are not identical by nature [49]. For example, the E-liquid vaporization process versus the tobacco combustion process result in exposure chambers with different physical environments such as temperature and humidity [40]. Furthermore, the volume of the human oral cavity, as determined by height, width and depth [50], is approximately 230 cm^3^, much smaller than the 2100 cm^3^ calculated for the exposure chambers used in this study [40]. This means that the effect of flavorless ECIG-aerosol or conventional cigarette smoke on oral commensal streptococci could be minimized in this investigation as compared to their effects *in vivo*. Alternatively, the *in vitro* results of this study could also be amplified when comparing the effect of aerosol and smoke on oral commensal streptococci. For example, our results clearly show that the effects of 50 puffs of conventional smoke strongly obliterates colony or biofilm formation in every oral species tested. However, it is important to note that our *in vitro* experiments do not have a mechanism for removal of aerosol or smoke materials as saliva would in the oral cavity *in vivo*, but rather are constantly exposed to these materials for the duration of the experiments.

Moreover, since our experimental design does not exhibit the properties of an open system, it cannot determine whether the overall effect of cigarette smoke on oral commensal bacteria is bacteriostatic or bactericidal. In the scenario where the smoke is bacteriostatic, such experimental design will allow for removal of potential bacteriostatic compounds and once the cigarette smoke materials fall below the minimal inhibitory concentration, the bacterial communities will be able to resume growth. It is important to note that the growth and/or architecture of biofilm communities may be altered at this point as a result of such cigarette smoke effects. Alternatively, if the smoke materials are bactericidal, the bacteria will be dead even after removal of such materials. This aspect deserves further study because the livelihood of the commensal bacteria is important to the homeostasis of the oral cavity, keeping it and the entire physiological system healthy.

## Conclusion

Our study indicates that flavorless ECIG aerosol (± nicotine) is less detrimental to the survival and growth of oral commensal streptococci than conventional cigarette smoke. This study opens the door for subsequent studies that could address the effect of flavorless ECIG aerosol on oral epithelial cells as well as the addition of flavoring agents to test all the above mentioned biological models.

## Acknowledgements

The authors gratefully acknowledge the technical expertise and assistance provided by Dr. Stan Kunigelis of the Center of Imaging and Analysis at Lincoln Memorial University (LMU) and Dr. Elizabeth McCain at Muhlenberg College (MC). The authors would also like to thank LMU and MC faculty (Drs. Gina DeFranco, Marten Edwards, Stacie Fairley, Douglas Fitzovitch, John Gibbons, Adam Gromley, Julie Hall, Amy Hark, Adam Rollins, Jordanna Sprayberry, Michael Wieting, Bruce Wightman and Jan Zieran) for graciously providing constructive criticism, comments and editorial assistance in the preparation of this manuscript.

## References

1. Glantz SA, Bareham DW. E-Cigarettes: Use, Effects on Smoking, Risks, and Policy Implications. Annu Rev Public Health. 2018;39: 215–235. doi:10.1146/annurev-publhealth-040617-013757

2. Hajek P, Etter J-F, Benowitz N, Eissenberg T, McRobbie H. Electronic cigarettes: review of use, content, safety, effects on smokers and potential for harm and benefit. Addiction. 2014;109: 1801–1810. doi:10.1111/add.12659

3. Jensen RP, Luo W, Pankow JF, Strongin RM, Peyton DH. Hidden formaldehyde in e-cigarette aerosols. N Engl J Med. 2015;372: 392–394. doi:10.1056/NEJMc1413069

4. Holliday R, Kist R, Bauld L. E-cigarette vapour is not inert and exposure can lead to cell damage. Evid Based Dent. 2016;17: 2–3. doi:10.1038/sj.ebd.6401143

5. Yu V, Rahimy M, Korrapati A, Xuan Y, Zou AE, Krishnan AR, et al. Electronic cigarettes induce DNA strand breaks and cell death independently of nicotine in cell lines. Oral Oncol. 2016;52: 58–65. doi:10.1016/j.oraloncology.2015.10.018

6. Muthumalage T, Prinz M, Ansah KO, Gerloff J, Sundar IK, Rahman I. Inflammatory and Oxidative Responses Induced by Exposure to Commonly Used e-Cigarette Flavoring Chemicals and Flavored e-Liquids without Nicotine. Front Physiol. 2017;8: 1130. doi:10.3389/fphys.2017.01130

7. Bahl V, Lin S, Xu N, Davis B, Wang Y, Talbot P. Comparison of electronic cigarette refill fluid cytotoxicity using embryonic and adult models. Reprod Toxicol. 2012;34: 529–537. doi:10.1016/j.reprotox.2012.08.001

8. Leigh NJ, Tran PL, O’Connor RJ, Goniewicz ML. Cytotoxic effects of heated tobacco products (HTP) on human bronchial epithelial cells. Tobacco Control. 2018;27: s26–s29. doi:10.1136/tobaccocontrol-2018-054317

9. Leigh NJ, Lawton RI, Hershberger PA, Goniewicz ML. Flavourings significantly affect inhalation toxicity of aerosol generated from electronic nicotine delivery systems (ENDS). Tob Control. 2016;25: ii81–ii87. doi:10.1136/tobaccocontrol-2016-053205

10. Sundar IK, Javed F, Romanos GE, Rahman I. E-cigarettes and flavorings induce inflammatory and pro-senescence responses in oral epithelial cells and periodontal fibroblasts. Oncotarget. 2016;7: 77196–77204. doi:10.18632/oncotarget.12857

11. Burne RA. Oral Streptococci… Products of Their Environment. J Dent Res. 1998;77: 445–452. doi:10.1177/00220345980770030301

12. Rosan B, Lamont RJ. Dental plaque formation. Microbes Infect. 2000;2: 1599–1607.

13. Kolenbrander PE. Oral Microbial Communities: Biofilms, Interactions, and Genetic Systems. Annual Review of Microbiology. 2000;54: 413–437. doi:10.1146/annurev.micro.54.1.413

14. Hajishengallis G, Lamont RJ. Beyond the red complex and into more complexity: the polymicrobial synergy and dysbiosis (PSD) model of periodontal disease etiology. Mol Oral Microbiol. 2012;27: 409–419. doi:10.1111/j.2041-1014.2012.00663.x

15. Aas JA, Paster BJ, Stokes LN, Olsen I, Dewhirst FE. Defining the Normal Bacterial Flora of the Oral Cavity. J Clin Microbiol. 2005;43: 5721–5732. doi:10.1128/JCM.43.11.5721-5732.2005

16. Colombo AV, da Silva CM, Haffajee A, Colombo APV. Identification of intracellular oral species within human crevicular epithelial cells from subjects with chronic periodontitis by fluorescence in situ hybridization. J Periodont Res. 2007;42: 236–243. doi:10.1111/j.1600-0765.2006.00938.x

17. Garnier F, Gerbaud G, Courvalin P, Galimand M. Identification of clinically relevant viridans group streptococci to the species level by PCR. J Clin Microbiol. 1997;35: 2337–2341.

18. Kolenbrander PE, Andersen RN, Blehert DS, Egland PG, Foster JS, Palmer RJ. Communication among oral bacteria. Microbiol Mol Biol Rev. 2002;66: 486–505, table of contents.

19. Kolenbrander PE, Egland PG, Diaz PI, Palmer RJ. Genome–genome interactions: bacterial communities in initial dental plaque. Trends in Microbiology. 2005;13: 11–15. doi:10.1016/j.tim.2004.11.005

20. Teles FR, Teles RP, Uzel NG, Song XQ, Torresyap G, Socransky SS, et al. Early microbial succession in re-developing dental biofilms in periodontal health and disease. J Periodontal Res. 2012;47: 95–104. doi:10.1111/j.1600-0765.2011.01409.x

21. Socransky SS, Haffajee AD, Cugini MA, Smith C, Kent RL. Microbial complexes in subgingival plaque. J Clin Periodontol. 1998;25: 134–144.

22. Heller D, Helmerhorst EJ, Gower AC, Siqueira WL, Paster BJ, Oppenheim FG. Microbial Diversity in the Early In Vivo-Formed Dental Biofilm. Appl Environ Microbiol. 2016;82: 1881–1888. doi:10.1128/AEM.03984-15

23. Hasegawa Y, Mans JJ, Mao S, Lopez MC, Baker HV, Handfield M, et al. Gingival epithelial cell transcriptional responses to commensal and opportunistic oral microbial species. Infect Immun. 2007;75: 2540–2547. doi:10.1128/IAI.01957-06

24. Thurnheer T, Belibasakis GN. Streptococcus oralis maintains homeostasis in oral biofilms by antagonizing the cariogenic pathogen Streptococcus mutans. Mol Oral Microbiol. 2018;33: 234–239. doi:10.1111/omi.12216

25. Gross EL, Beall CJ, Kutsch SR, Firestone ND, Leys EJ, Griffen AL. Beyond Streptococcus mutans: dental caries onset linked to multiple species by 16S rRNA community analysis. PLoS ONE. 2012;7: e47722. doi:10.1371/journal.pone.0047722

26. Huang X, Browngardt CM, Jiang M, Ahn S-J, Burne RA, Nascimento MM. Diversity in Antagonistic Interactions between Commensal Oral Streptococci and Streptococcus mutans. Caries Res. 2018;52: 88–101. doi:10.1159/000479091

27. Liu Y, Palmer SR, Chang H, Combs AN, Burne RA, Koo H. Differential oxidative stress tolerance of Streptococcus mutans isolates affects competition in an ecological mixed-species biofilm model. Environ Microbiol Rep. 2018;10: 12–22. doi:10.1111/1758-2229.12600

28. Herrero ER, Slomka V, Bernaerts K, Boon N, Hernandez-Sanabria E, Passoni BB, et al. Antimicrobial effects of commensal oral species are regulated by environmental factors. J Dent. 2016;47: 23–33. doi:10.1016/j.jdent.2016.02.007

29. Holmlund A, Lampa E, Lind L. Oral health and cardiovascular disease risk in a cohort of periodontitis patients. Atherosclerosis. 2017;262: 101–106. doi:10.1016/j.atherosclerosis.2017.05.009

30. Li M, Huang R, Zhou X, Zhang K, Zheng X, Gregory RL. Effect of nicotine on dual-species biofilms of Streptococcus mutans and Streptococcus sanguinis. FEMS Microbiol Lett. 2014;350: 125–132. doi:10.1111/1574-6968.12317

31. Huang R, Li M, Ye M, Yang K, Xu X, Gregory RL. Effects of Nicotine on Streptococcus gordonii Growth, Biofilm Formation, and Cell Aggregation. Appl Environ Microbiol. 2014;80: 7212–7218. doi:10.1128/AEM.02395-14

32. Tomar SL, Asma S. Smoking-Attributable Periodontitis in the United States: Findings From NHANES III. Journal of Periodontology. 2000;71: 743–751. doi:10.1902/jop.2000.71.5.743

33. Bagaitkar J, Daep CA, Patel CK, Renaud DE, Demuth DR, Scott DA. Tobacco smoke augments Porphyromonas gingivalis-Streptococcus gordonii biofilm formation. PLoS ONE. 2011;6: e27386. doi:10.1371/journal.pone.0027386

34. Moon J-H, Lee J-H, Lee J-Y. Subgingival microbiome in smokers and non-smokers in Korean chronic periodontitis patients. Mol Oral Microbiol. 2015;30: 227–241. doi:10.1111/omi.12086

35. Calafat AM. Determination of tar, nicotine, and carbon monoxide yields in the mainstream smoke of selected international cigarettes. Tobacco Control. 2004;13: 45–51. doi:10.1136/tc.2003.003673

36. Cobb E, Hall J, Palazzolo DL. Induction of Metallothionein Expression After Exposure to Conventional Cigarette Smoke but Not Electronic Cigarette (ECIG)-Generated Aerosol in Caenorhabditis elegans. Front Physiol. 2018;9: 426. doi:10.3389/fphys.2018.00426

37. Palazzolo DL, Nelson JM, Ely EA, Crow AP, Distin J, Kunigelis SC. The Effects of Electronic Cigarette (ECIG)-Generated Aerosol and Conventional Cigarette Smoke on the Mucociliary Transport Velocity (MTV) Using the Bullfrog (R. catesbiana) Palate Paradigm. Front Physiol. 2017;8: 1023. doi:10.3389/fphys.2017.01023

38. Trehy ML, Ye W, Hadwiger ME, Moore TW, Allgire JF, Woodruff JT, et al. Analysis of Electronic Cigarette Cartridges, Refill Solutions, and Smoke for Nicotine and Nicotine Related Impurities. Journal of Liquid Chromatography & Related Technologies. 2011;34: 1442–1458. doi:10.1080/10826076.2011.572213

39. Murray J. Nicotine and what else?: HPLC elution optimization for the analysis of alkaloids found in electronic cigarettes. Honors Theses. 2014; Available: https://scholar.utc.edu/honors-theses/3

40. Palazzolo DL, Crow AP, Nelson JM, Johnson RA. Trace Metals Derived from Electronic Cigarette (ECIG) Generated Aerosol: Potential Problem of ECIG Devices That Contain Nickel. Front Physiol. 2017;7. doi:10.3389/fphys.2016.00663

41. Sutton S. Accuracy of plate counts. Journal of validation technology. 2011;17: 42–46.

42. Kolenbrander PE. Multispecies communities: interspecies interactions influence growth on saliva as sole nutritional source. Int J Oral Sci. 2011;3: 49–54. doi:10.4248/IJOS11025

43. Shah SA, Ganesan SM, Varadharaj S, Dabdoub SM, Walters JD, Kumar PS. The making of a miscreant: tobacco smoke and the creation of pathogen-rich biofilms. NPJ Biofilms Microbiomes. 2017;3: 26. doi:10.1038/s41522-017-0033-2

44. Zonuz AT, Rahmati A, Mortazavi H, Khashabi E, Farahani RMZ. Effect of cigarette smoke exposure on the growth of Streptococcus mutans and Streptococcus sanguis: an in vitro study. Nicotine Tob Res. 2008;10: 63–67. doi:10.1080/14622200701705035

45. Abiko Y, Sato T, Mayanagi G, Takahashi N. Profiling of subgingival plaque biofilm microflora from periodontally healthy subjects and from subjects with periodontitis using quantitative real-time PCR. J Periodont Res. 2010;45: 389–395. doi:10.1111/j.1600-0765.2009.01250.x

46. Genco CA, Van Dyke T, Amar S. Animal models for Porphyromonas gingivalis-mediated periodontal disease. Trends Microbiol. 1998;6: 444–449.

47. Olsen I, Progulske-Fox A. Invasion of Porphyromonas gingivalis strains into vascular cells and tissue. J Oral Microbiol. 2015;7. doi:10.3402/jom.v7.28788

48. Pussinen PJ, Tuomisto K, Jousilahti P, Havulinna AS, Sundvall J, Salomaa V. Endotoxemia, immune response to periodontal pathogens, and systemic inflammation associate with incident cardiovascular disease events. Arterioscler Thromb Vasc Biol. 2007;27: 1433–1439. doi:10.1161/ATVBAHA.106.138743

49. Sosnowski TR, Odziomek M. Particle Size Dynamics: Toward a Better Understanding of Electronic Cigarette Aerosol Interactions With the Respiratory System. Front Physiol. 2018;9: 853. doi:10.3389/fphys.2018.00853

50. Kaufman JW, Farahmand K. In vivo measurements of human oral cavity heat and water vapor transport. Respir Physiol Neurobiol. 2006;150: 261–277. doi:10.1016/j.resp.2005.05.016

